# Merging conformational landscapes in a single consensus space with FlexConsensus algorithm

**DOI:** 10.1101/2025.01.07.631840

**Authors:** D. Herreros, C.P. Mata, C.O.S. Sorzano, J.M. Carazo

## Abstract

The analysis of structural heterogeneity in CryoEM is experimenting a significant breakthrough toward estimating more accurate, richer, and interpretable conformational landscapes directly derived from experimental data. The increasing number of new methods designed to tackle the heterogeneity challenge reflects these new paradigms, allowing users to understand protein dynamics better. However, a question remains on how the estimation of different heterogeneity algorithms compare, which is essential to properly determine the reliability, stability, and correctness of the conformational landscapes and the structural transitions extracted from the CryoEM images. The intrinsic differences arising from the approximation and implementation of every heterogeneity algorithm make the definition of a consensus complex, a problem that remains unsolved. To overcome the challenges of comparing heterogeneity algorithms, we introduce our new FlexConsenus algorithm in this work. FlexConsensus relies on a multi-autoencoder architecture to learn the commonalities and differences of several conformational landscapes, allowing one to place them in a shared consensus space with enhanced reliability. In addition, FlexConsensus enables the derivation of error metrics from the consensus space to measure the reproducibility of the heterogeneity estimations. Thanks to the previous consensus metrics, it is possible to clean the original particle datasets further based on the differences in the estimation of their structural variability among different algorithms and/or different runs, allowing users to focus their analysis only on those particles with a stable estimation of their structural variability.

## Introduction

Cryo-electron microscopy (Cryo-EM) is experiencing a paradigm change in exploring conformational variability from experimental data. Unlike classical 3D classification algorithms (1) that are confined to a set of reduced and stable states, new heterogeneity algorithms focus on estimating richer and more complete conformational landscapes with the possibility of retrieving any conformation from the approximated continuum.

Heterogeneity algorithms may be classified according to the information estimated from the Cryo-EM data. Heterogeneous reconstruction methods (2; 3; 4; 12) rely on directly estimating electron density maps from a continuous function that maps every point in the conformational landscape to a 3D volume. On the other hand, deformation field-based methods directly estimate the motions responsible for driving a reference state to a new conformational state represented in a particle dataset (7; 6; 5).

The variety of methods, approaches, and implementations allows one to explore the structural variability of any given dataset through a systematic approach. However, the large pool of algorithms also introduces the challenge of comparing different results. In addition, the accuracy of the current heterogeneity analysis methods is still being studied. Because of that, the extraction of consensus solutions among different approaches is precious.

To overcome the challenge of comparing conformational landscapes obtained from different methods and/or different runs, we propose in this work a new deep learning algorithm called FlexConsensus. FlexConsensus introduces a multi-autoencoder architecture designed to merge different conformational landscapes into a common latent space posteriorly decoded back into the original representations input to the network. In this way, it is possible not only to analyze the common conformational space but also to determine a consensus metric that measures the stability of every estimation in the original conformational landscapes to filter out only those regions with higher confidence. Lastly, the method also allows for converting among different conformational landscape representations, simplifying the comparison of techniques.

## Results

The following sections present some use cases to evaluate and discuss the precision and performance of FlexConsensus under different conditions. The first two proposed cases define a synthetic and controlled dataset to better understand the mode of operation of the new consensus method and the results expected from this type of analysis. Following these two cases, the new process is tested with two experimental datasets to assess the behavior of the analysis in more realistic scenarios.

We remark here that FlexConsensus learns by default a consensus space with three dimensions, although the latent space dimensionality is exposed as a customizable parameter in the corresponding Scipion (?) protocol form. All the consensus landscapes presented in the following sections were defined to have three dimensions as set by default.

The workflow to train the network is described in detail in the Methods sections and in Figure 8.

### Simulated conformational landscape with different dimensions

One of the first issues when comparing different heterogeneity algorithms is the variable nature of the conformational landscapes they generate. Apart from the intrinsic differences related to the assumptions that they consider, a mismatch may also exist in their dimensions. The dimensional differences make the consensus evaluation challenging and should be addressed in the first instance to yield meaningful and interpretable comparison metrics.

In FlexConsensus, we propose a multi-encoder architecture relying on a common latent space to overcome the previous issue, as further described in the *Methods* section. The proposed architecture effectively reduces the multidimensional conformational landscapes of different methods to a new latent space with unified dimensions, thus simplifying their comparison. Assuming that the number of input cryoEM images is N and the number of different input landscapes is M, the number of points in the common latent space will be NxM, and we will always know from which of the input landscapes a particular point in the common latent space is coming from. Note that each of the input conformational spaces to FlexConsensus may be a latent space in its own right (because it may come from some form of auto-encoder architecture, for example); however, we will always refer to them as input spaces, keeping the reference to latent space to the new and common space obtained by FlexConsensus

To determine the performance of our multi-encoder architecture under the previously described conditions, we designed a simple synthetic case with two different input landscapes. We will refer to “Consensus space 1” and “Consensus space 2” as the mapping into the new common latent space of the points coming from the first and second input spaces, respectively. It should be noted that FlexConsensus does not restrict the number of conformational landscapes to be analyzed, and the choice of two input spaces is only for simplicity in the presentation.

The synthetic landscapes were generated using sine and cosine functions for the even and odd dimensions of the target space. The generation process is thus similar to the definition of the well-known Lissajous curves but applied to the multi-dimensional case. This process allows the generation of spaces with similar characteristics and varying dimensions. We can express the equation of the curves as:

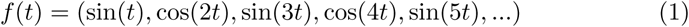

where *t* takes values in [0, 2*π*].

For the current tests, we generated two input spaces of 1,000 points in the range [0, 2*π*], each with 8 and 39 dimensions, respectively, replicating the default number of dimensions of two entirely different heterogeneity algorithms in the field: CryoDRGN (2) and Zernike3D (7).

The results obtained using FlexConsensus with the previous synthetic spaces are summarized in Figure 1. The training workflow starts with the input spaces being forwarded through their encoding network (as further detailed in the Methods section), yielding a consensus latent space composed of “Consensus space 1” and “Consensus space 2”. The consensus space is then forwarded through the decoders responsible for recovering the original input spaces from the consensus latent space. At this point, we can measure a representation loss function between the decoded and the input spaces, which will be used by the network in conjunction with other losses during the training phase.

**Figure 1:**
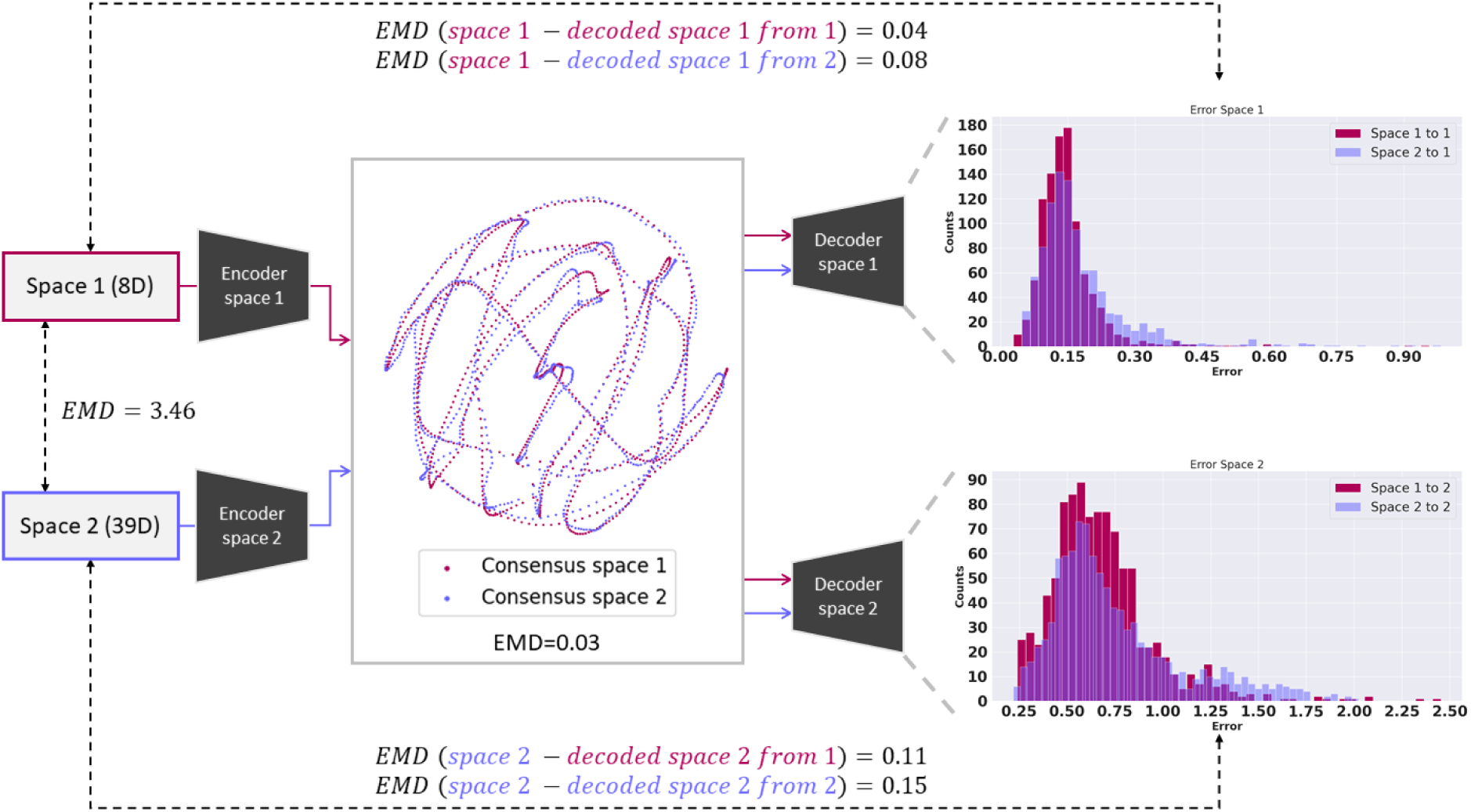
FlexConsensusanalysis resulting from the comparison of two synthetic spaces with a varying number of e resulting consensus space, error histograms derived from the representation error computed by decoded and input spaces, and the EMD evaluation metrics estimated from the analysis are also Figure. The analysis shows that FlexConsensus identified that the two input spaces have similar which is to be expected by construction. Apart from having a similar consensus space for both ysis of the error metrics resulting from comparing the input and decoded spaces is low, showing the cy even when faced with multidimensional datasets.

The differences in the dimensions between the input spaces make it hard to determine whether the network has produced an appropriate consensus space. Therefore, we proposed two ways to evaluate the network’s accuracy: analyzing 1) the final representation loss computed as the distance between the input and decoded spaces and 2) the Earth Mover’s Distance (EMD) between the distance distributions among different spaces. It is considered that distances effectively capture the spatial characteristics of a given space, making the EMD an excellent candidate for determining the accuracy of the network predictions independently of their initial dimension.

The results obtained with the synthetic dataset proposed before are summarized in Figure 1. The analysis starts by measuring the EMD between the input spaces to have a reference value to compare the following EMD metrics to be measured. Even if the input spaces have similar shapes by constructions, the EMD results in a large value, mainly due to the differences in the dimensions of the inputs.

The next step consists of encoding the input spaces into the common latent space, leading to “Consensus space 1” and “Consensus space 2” as shown in Figure 1. Similar to the case before, the EMD between the “Consensus space 1” and “Consensus space 2” was measured, yielding a small value compared to the reference EMD computed before. Both the EMD and the visual inspection of the landscapes effectively show that FlexConsensus has correctly determined that the input spaces were very similar, as expected (note that the curves of red and blue points in the consensus landscape map are virtually one on top of the other).

Lastly, “Consensus space 1” and “Consensus space 2” were forwarded through the different decoders, generating four decoded spaces. Two of these spaces correspond to the transformation of “Consensus space 1” and “Consensus space 2” into the first input space, while the other two come from the transformation of “Consensus space 1” and “Consensus space 2” into the second input space. At this point, we proposed two evaluation metrics: the first one is the EMD between the input spaces and their analogous decoded spaces, yielding a total of 4 measurements. Again, these metrics show a small value as FlexConsensus has correctly learned to reproduce the input spaces from the consensus latent space accurately.

The second evaluation is obtained from the pairwise comparison of the input spaces and their analogous decoded spaces (corresponding to the representation loss used to train the neural network), represented as a histogram to simplify their understanding. In agreement with the EMD metrics, the histograms presented in Figure 1 show that the representation error is small, meaning that FlexConsensus has correctly learned to decode the input spaces from the consensus.

### Simulated conformational landscape with variable stability

Although the dataset analyzed in the previous section helps evaluate the method’s performance when faced with multidimensional data sets, it fails to reproduce the errors that disrupt the conformational landscapes in a real-case scenario. Among the errors that affect the estimation of conformational landscapes, the noise contaminating the experimental cryoEM images is the most predominant. This noise is transmitted to conformational landscape estimations, which is one of the reasons why consensus among several estimations is desired to improve the reliability and interpretation of conformational landscapes.

Therefore, a new test case is proposed to determine FlexConsensus’ ability to discern which regions of the input spaces can be trusted and which areas are more questionable due to estimation differences. Similarly to the previous experiment, we generated two input spaces following the Lissajous-like construction method proposed earlier. However, in this case, constructing the input spaces includes some additional steps to simulate some of the effects experimental latent spaces may have.

Focusing on the first input space, the construction starts with generating 1,000 8D points, as done in the previous section. These 1,000 points are then duplicated, followed by a translation of the duplicated points so they do not overlap with the original ones. This way, we obtained two replicas of the same 8D space at distinct locations. Lastly, gaussian noise was applied to one of these replicas, leading to the final input space consisting of one structured region following the Lissajous curves (the original 1,000 points) and another region that did not show any apparent structure due to the added high noise (the noisy duplicated 1,000 points). The standard deviation of the noise added to the points was large enough to completely disrupt their structure.

The second input space follows the same construction process as the previous one, starting with 1,000 39D points. In addition, this second space was also rotated by 90 degrees around the center of mass of the complete space. This rotation simulates the randomness in the orientation of experimental spaces, as they could be randomly rotated even if the method responsible for generating that space is executed twice with the same parameters.

The new input spaces (two input spaces, each with 1000 noise-free points and 1000 noisy data points) were fed to FlexConsenus to train the neural network. This allowed the method to learn the common consensus space and decode the input spaces directly to evaluate the representation losses. Note that we will have 2×2000 points in the consensus latent espace, 2000 from each input space. However, and only for the sake of simplicity in the presentation, we will split the analysis of FlexConsensus into two sections. In the first section, we will address the performance holding the two noise-free spaces (Figure 2a)), and in the second one, we will study the results with the noisy spaces (Figure 2b). Similarly to the lay-out followed in Figure 1, the analysis starts by measuring the EMD between the points in the input spaces to get a reference value to compare, it follows with a visual inspection of the consensus latent space, and it ends with the presentation of error metrics in the form of histograms (see Methods section) and the EMD between the two decoded spaces and the input spaces.

**Figure 2:**
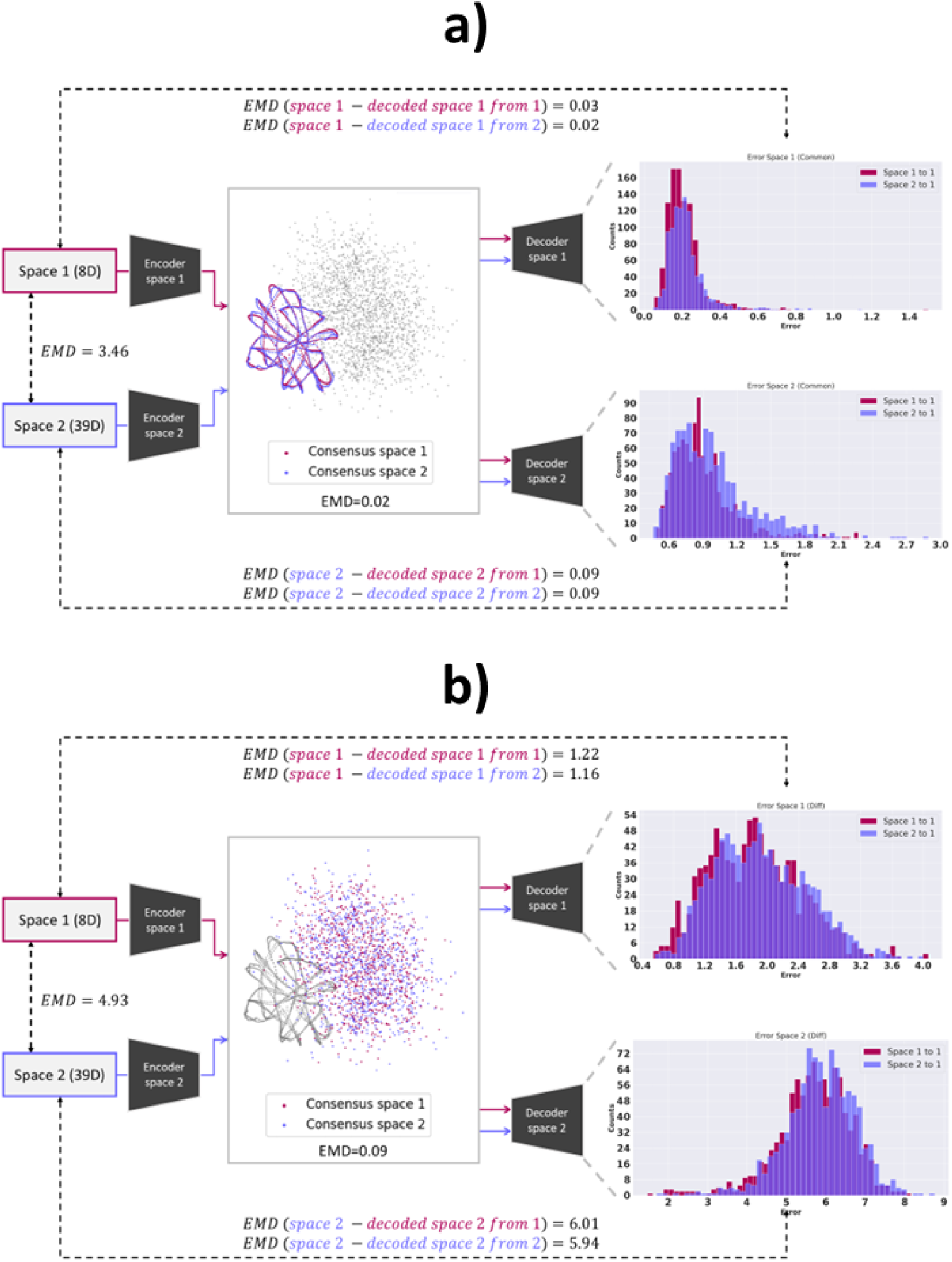
FlexConsensus analysis resulting from the comparison of two synthetic spaces with a varying number of addition, the synthetic spaces also mimic the presence of noise and other errors found in typical ent spaces. Panel a) compares the consensus space corresponding to those points in the input spaces ded to them. The resulting consensus landscape, error histograms derived from the representation by comparing the decoded and input spaces, and the EMD evaluation metrics estimated from the presented in the Figure. These results show that FlexConsensus successfully identified these regions uctural properties. In contrast, Panel b) focuses the analysis on the points affected by noise in the he analysis shows that FlexConsensus identifies these two regions as unreliably estimated, as they

Regarding the analysis of the 2×1,000 noiseless structured points in the input spaces, results are summarized in Figure 2a. As expected, the EMD value between input spaces is the same as shown in Figure 1, as the noiseless points were constructed following the same process in both experiments. As for the evaluation of the noiseless “Consensus space 1” and “Consensus space 2” (blue and red points in Figure 2a), visual inspection shows that the two sets of 1000 points map virtually map one on top of each other; note that in this consensus space, the 2×2000 points coming from the input spaces are shown, although only the 2×1000 points corresponding to the noiseless input spaces are colored (the additional 2×1000 points coming from the noisy spaces are kept in gray and will be analyzed in the second section). Lastly, “Consensus space 1” and “Consensus space 2” were fed to the decoders to generate the four decoded spaces, similarly to the process followed with the first test case. Therefore, it is possible to measure the EMD between the two decoded spaces analogous to the first input space, the EMD between the two decoded spaces analogous to the second input space, and the histograms computed from the representation errors arising from the pairwise comparison of the input and the decoded spaces. As can be seen from the histograms and the small EMD distances, FlexConsenus has effectively learned to reproduce the noiseless points in the input spaces from the noiseless points in the common consensus space.

The second section refers to analyzing the two very noisy input spaces, each with 1000 points, as shown in Figure 2b (blue and red points). Again, we start by measuring the EMD between the noisy points in the input space, leading to a larger reference value due to the addition of Gaussian noise. The analysis by visual inspection of the noisy “Consensus space 1” and “Consensus space 2” shows two clouds of points without structure or overlap. Additionally, the EMD measured between them is very large, indicating that FlexConsensus fails to identify any similarity between these points. This is expected, as FlexConsensus should be able to determine that the two sets of very noisy points in the two input spaces are no longer similar due to the noise. Lastly, the analysis of the four spaces decoded from the noisy “Consensus space 1” and “Consensus space 2” show a similar trend, leading to larger errors in both the EMD measurements and the histograms obtained from the representation errors coming from the comparison with the 1000 noisy points in the input spaces.

The previous experiment shows that FlexConsensus is sensitive enough to correctly determine which regions in different latent spaces are more or less reliable, even if the original spaces have different characteristics, orientations, or dimensions.

### Consensus results on the EMPIAR 10028 dataset

The next step in evaluating FlexConsensus’s capabilities is applying the method to a more realistic scenario. Therefore, we proposed the evaluation of FlexConsensus with the EMPIAR 10028 dataset (8), a well-known and well-studied dataset showing different conformational states of the *P. falciparum* 80S ribosome bound to emetine. This data set has been widely applied as a test case for the most recently developed heterogeneity methods, yielding a conformational landscape with well-defined features. In addition, the experimental images in the dataset mainly capture continuous conformational changes, although there is also a compositional variability component. Thanks to all these characteristics, the EMPIAR 10028 dataset supposes a realistic yet controlled scenario to evaluate the consensus when considering conformational landscapes estimated by different methods.

The dataset was first preprocessed with CryoSPARC (9) inside Scipion (10) to yield a set of appropriately characterized experimental images to be analyzed by heterogeneity algorithms. This involves estimating CTF information and particle alignments, which were subjected to a consensus analysis to improve their stability (11).

A set of 50k particles was obtained from the preprocessing step and further analyzed for conformational variability. The study was carried out with two different methods: HetSIREN (12) and Zernike3D (7). These two methods follow very different approaches to estimating conformational variability, HetSIREN being a heterogeneous reconstruction/refinement algorithm and Zernike3D being a deformation field-based method. Therefore, HetSIREN can extract continuous and compositional variability, while Zernike3D focuses on extracting continuous motions. Moreover, each method defines a conformational landscape with different dimensions, similar to the synthetic datasets analyzed in the previous sections.

In total, three independent conformational landscapes were estimated, corresponding to the execution of HetSIREN in reconstruction mode, the execution of HetSIREN in refinement mode, and the execution of Zernike3D. The main difference between the reconstruction modes of HetSIREN is the algorithm’s initialization: in reconstruction mode, the initial volume required by HetSIREN is initialized to have only zero values. In contrast, HetSIREN in refinement mode receives as the initial volume a map calculated from all initial images. These landscapes were then fed to FlexConsensus to generate the consensus landscape and the error metrics needed to determine the reliability of the three estimations.

It is important to mention that FlexConsensus also works with estimations obtained from different executions of the same method under the same conditions, allowing the study of the consistency and reliability of a single algorithm’s estimations. Although we could have executed HetSIREN in this way, we considered comparing this method’s two modes of operation to determine if they impact the estimated conformational landscapes.

Before analyzing the results obtained for this dataset, we include some terminology that will be used from now on when comparing different results. We will refer to the isolines computed from the density distribution of the points in the consensus space associated with method X as “X contour.” These isolines will help identify the location of the points coming from input space X in the common consensus space, allowing us to compare the differences among the different methods in the consensus quickly.

The results obtained from the analysis of the common consensus space learned by FlexConsenus are summarized in Figure 3. Panel a) presents the common space with the previously described contours associated with each input space. Atop panel a), the “Zernike3D contour” is presented, showing only those points coming from the Zernike 3D landscape (i.e., the mapping onto the common space of the point cloud corresponding to the Zernike3D analysis). At the bottom, a similar representation is followed with the points coming from the two HetSIREN landscapes. The general disposition of the three contours is visually similar, indicating that the strategy of mapping all input results to the same common space is adequate. In detail, a difference between Zernike3D and HetSIREN is noticeable in the common landscape at the top rightmost corner, and it is highlighted by a circle drawn with a broken green line. The Zernike3D contour in this area is unstructured (points are sparse), unlike both HetSIREN landscapes that show a highly ordered contour. Interestingly, the analysis of the maps coming from these points indicates that they correspond to specimens presenting a substantial compositional variation (they lack the 40S subunit, showing only the 70S one), which Zernike3D, by design, could not capture. The point clouds within the purple and yellow broken line circles correspond to particles presenting internal motions, which all three methods have captured well. In addition to the results displayed in panel a), we include in panel b) a highlight of the regions estimated to be less reliable for every method according to FlexConsensus. In Zernike3D, the higher errors are associated with the compositional components, which agrees with the results previously discussed. In the case of HetSIREN, these regions are found in the periphery of the landscape, which are also regions with a lower density than the central cloud. These differences are further highlighted in Supplementary Figure 1. It should be noted that the consensus landscape has three dimensions as specified when defining the network, although Figure 3 only shows a 2D projection. Thus, some of the highest error points in HetSIREN seem to be located within the landscape, although the visualization in 3D reveals that they are also located in the periphery.

**Figure 3:**
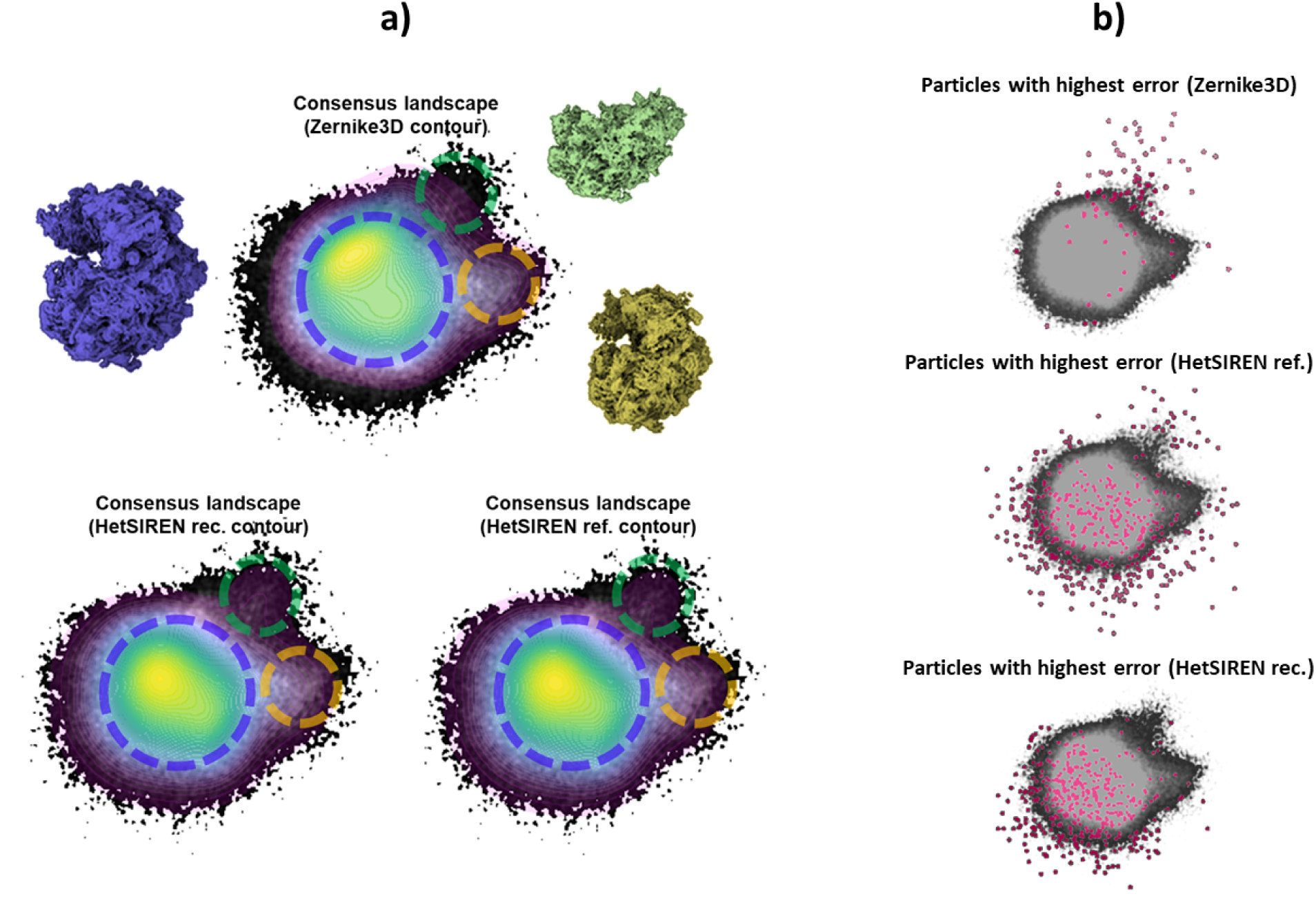
Consensus landscapes obtained by FlexConsensus for the EMPIAR 10028 dataset (8). Three conforwere input to the FlexConsensus analysis, obtained from three independent runs of the following REN in reconstruction/refinement mode (12) and Zernike3D (7). Panel a) compares the overall cone against each consensus landscape generated from each input (shown as a contour representation). show a region identified by both HetSIREN runs that is not present in Zernike3D, corresponding changes of the ribosome. It should be noted that, due to the approximation of each method, Hettify compositional variability while Zernike3D focuses on continuous conformational changes. Panel e regions assigned a higher consensus error for each method. For Zernike3D, the highlighted regions he region where the biggest conformational variability change has been detected by HetSIREN, d as Zernike3D cannot properly identify compositional variability. In the case of HetSIREN, these ted on the border regions of the 3D consensus landscape that are further from the main cloud.

To further support the results observed from the inspection of the consensus space, in Figure 4, we show the error histograms computed from the representation error obtained from the comparison of the input spaces against their analogous decoded spaces. Similarly to the previous sections, each consensus space (i.e., the points in the consensus space coming from the Zernike3D and HetSIREN landscapes) can be decoded towards each input space. Therefore, we have three decoded spaces analogous to the Zernike3D, three to HetSIREN in reconstruction mode, and three to HetSIREN in refinement mode.

**Figure 4:**
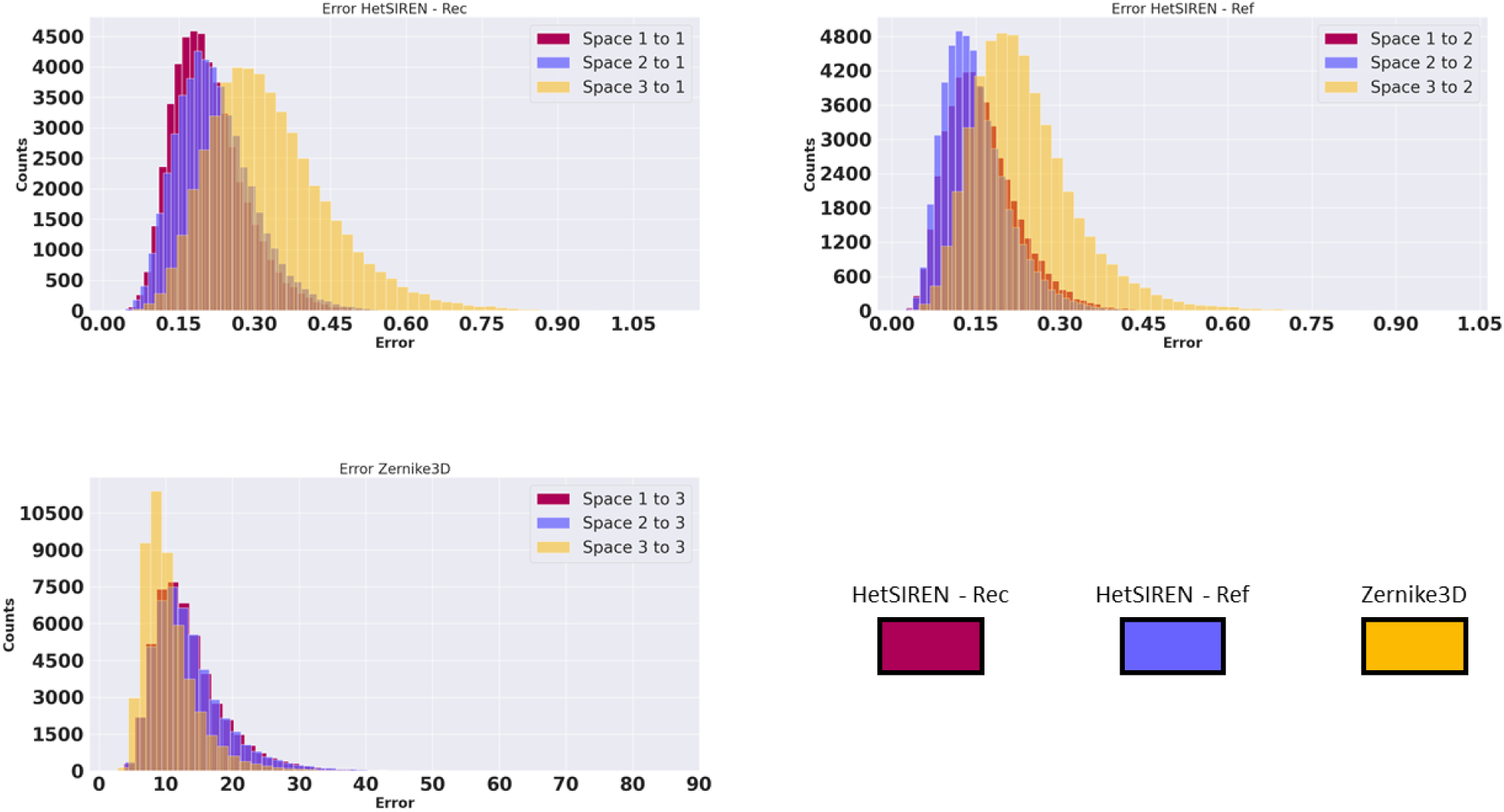
FlexConsensus error histograms computed for the EMPIAR 10028 dataset. The error metric corresponds oot Mean Squared Error between every original space and the spaces decoded by FlexConsensus sus space. The three images presented show a similar error tendency for all the decoders: the two utions remain more similar, meaning that the two independent estimations have similar characterisable for most particles. In contrast, Zernike3D deviates more from the HetSIREN, an effect mainly o the differences in the approximation followed by each method and the differences in the type of can estimate. Apart from these differences, one can see that the distribution of the errors is similar s, and it allows us to easily identify and exclude those particles on the right tail of the distributions, nes estimated with lower reliability based on these three executions.

The histograms show a similar distribution of the errors independently on the landscape selected for the comparison. As expected, the two HetSIREN runs are more similar, meaning the two executions were relatively stable. This also means that the HetSIREN operation mode has a non-significant effect in determining the conformational landscape in this specific case. It is also possible to detect how Zernike3D errors tend to deviate more from HetSIREN and the effect arising from the differences in the type of heterogeneity both methods can detect. From the histograms, one can quickly identify and interactively determine the particles to exclude from future analysis, as they accurately summarize how reliable the heterogeneity estimation was for every particle.

### Consensus results on SARS-CoV-2 D614G dataset

To further assess FlexConsensus’s capabilities under different experimental conditions, we evaluated the method with the SARS-CoV-2 D614G spike using a data set obtained at Prof. Subramanian laboratory (U. of Seattle) and currently under study (James Krieger, personal communication). This protein is well-characterized and exhibits a wide range of motions, primarily affecting, in the prefussion state, the Receptor Binding Domains (RBD) and the N-terminal domains (NTD). The RBD transitions are particularly interesting among the motions the spike undergoes due to their localized and highly dynamic nature. Similarly to the pipeline followed in the previous section, the experimental data set was preprocessed within Scipion (10), leading to 440k particle images with CTF and angular information. The structural variability captured by the particles was then approximated with two different software: HetSIREN (12) and CryoDRGN (2). In contrast to Zernike3D discussed in the previous section, both HetSIREN and CryoDRGN follow the heterogeneous reconstruction approximation to extract the conformational landscape from a set of images. Their conformational landscapes should be more comparable since they follow a similar approximation to solve the structural heterogeneity problem.

Again, the analysis will follow the same terminology introduced in the previous experiment: “X contour,” the isolines determining the location of the points coming from input space X in the common consensus space.

The reliability of the two independently estimated landscapes was analyzed by FlexConsensus, leading to the consensus landscapes presented in Figure 5. Panel a) includes the common consensus landscape and the HetSIREN and Cry-oDRGN contours. An initial inspection of these representations reveals three central structural regions corresponding to the RBDs in three down, one up, and two up states correctly identified by both methods. However, HetSIREN concentrates more on the three-down and one-up states, unlike CryoDRGN, which focuses more on the two-up and less on the three-down. In addition, panel b) shows the location of the images estimated to have a more significant consensus error based on the structural variability estimation from HetSIREN and CryoDRGN. In both cases, these particles are located in the periphery of the consensus landscape, this arrangement being a more prominent feature in the case of HetSIREN. Again, it is important to remark that the consensus landscape lives in a 3D space. Thus, some of the highest error points shown in Figure 5b appear to be in the interior of the landscape. However, the visualization in 3D reveals that they are also located in the periphery. Particles located in the periphery are usually associated with states estimated wrongly due to bad particle images or random estimation errors. Overall, the presence of these periphery particles is not significant compared to the inner region of the consensus landscapes, where a more thorough analysis can be performed to properly describe if the differences in the distribution of state estimated by different methods are significant.

**Figure 5:**
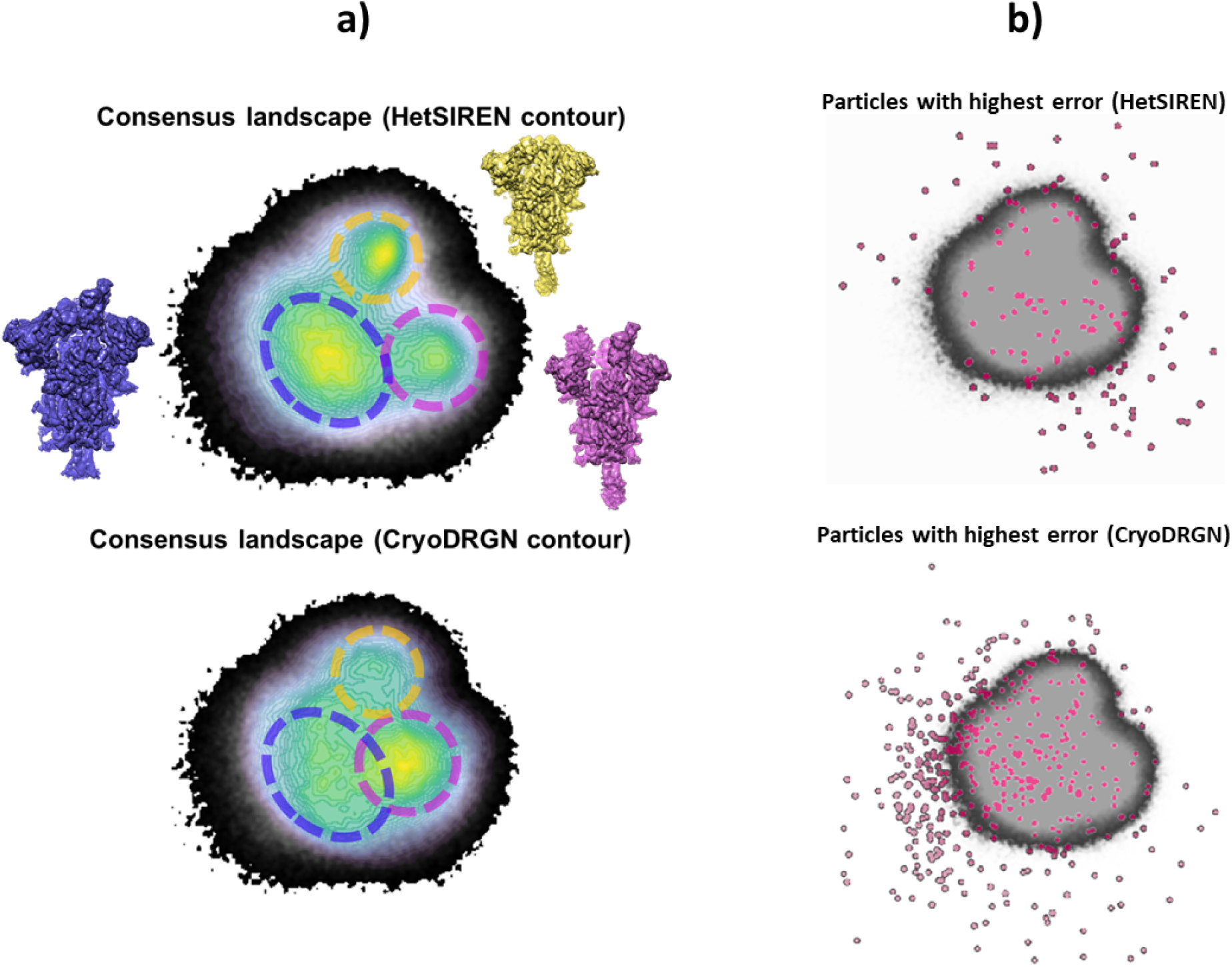
Consensus landscapes obtained by FlexConsensus for the SARS-CoV-2 D614G spike variant. Two conces were input to the FlexConsensus analysis, obtained from two independent runs of the following REN in reconstruction mode (12) and CryoDRGN (2). Panel a) compares the overall consensus st each consensus landscape generated from each input (shown as a contour representation). The capes show three main regions corresponding to three different conformational states of the spike ne up, and two up. Based on the consensus, it is possible to see that both methods correctly identify ural states. However, HetSIREN is more evenly distributed than CryoDRGN, which identifies two minently. Panel b) highlights the regions assigned a higher consensus error for each method. In both ost unstable particles tend to organize in the periphery of the consensus conformational landscape, regions of the landscape quite stable.

To more quantitatively assess the reliability of the structural states estimated by HetSIREN and CryoDRGN, we evaluated the representation error between the input spaces and those decoded from the consensus space. Four spaces were decoded, obtained when forwarding the HetSIREN and CryoDRGN consensus spaces through the decoders responsible for generating the original two spaces from the consensus. These four spaces were then used to compute the representation errors represented as histograms in Figure 6. The error histograms reveal that the heterogeneity estimation for most particles is stable, apart from a small fraction of the images estimated to have a more significant consensus error. This result agrees with the findings previously discussed directly from the consensus landscapes.

**Figure 6:**
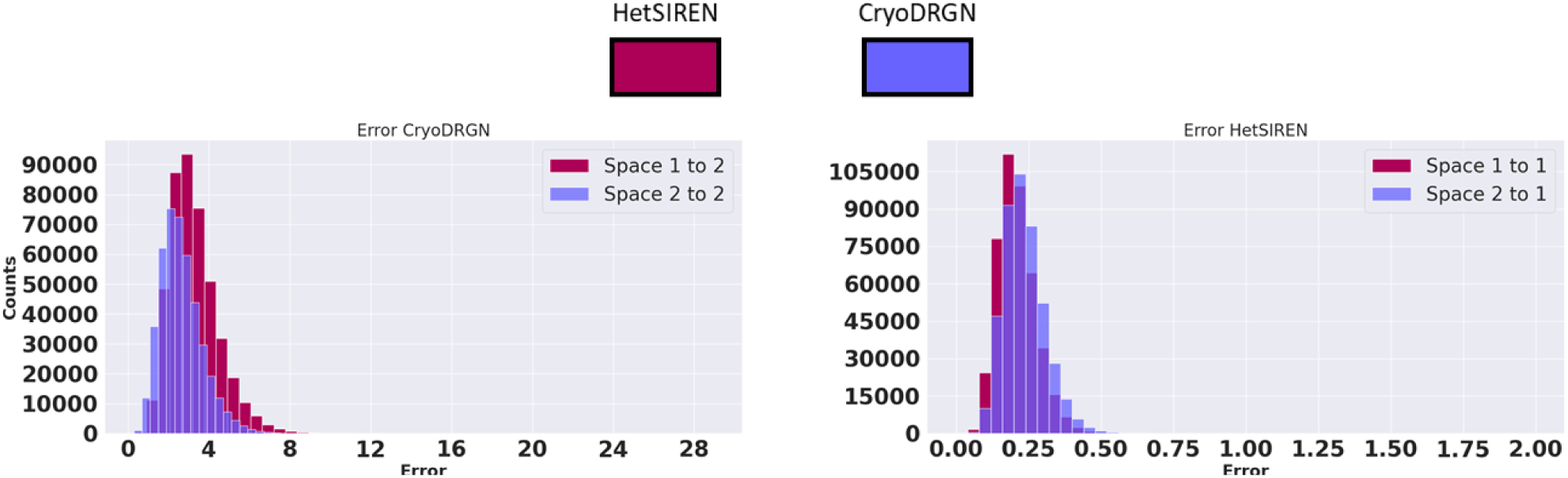
FlexConsensus error histograms computed for the SARS-CoV-2 D614G spike. The error metric corredard Root Mean Squared Error between every original space and the spaces decoded by FlexConconsensus space. The histograms reveal that the heterogeneity estimations yielded by HetSIREN are, in general terms, reasonably stable, apart from a few fractions of particles that are not reliably ing to their larger consensus error.

The shared consensus space and consensus metrics obtained from the previous analysis may also filter the landscape toward a stabilized representation with a more reliable state distribution. Based on the consensus, it is possible to define a statistical framework to determine a significant threshold based on the similarity of the distribution of states defined by the consensus landscapes. To that end, it is possible to work under the assumption that the distributions of states of different methods in the consensus space should be the same, allowing to derive the threshold that fulfills the previous assumption.

Following the previous reasoning, we proposed an approach based on Flex-Consensus to derive the previous threshold, whose application is summarized in Figure 7 for the SARS-CoV-2 D614G dataset under study. Thanks to this approach, it is possible to filter the original consensus space towards a new distribution of states that agrees with the estimation of the methods involved in the consensus (e.g. HetSIREN and CryoDRGN for the current case). This new representation of the consensus space allows us to analyze the distribution of states and its derived implications in a more reliable, significant, and accurate manner, compared to the estimation of a single method alone.

**Figure 7:**
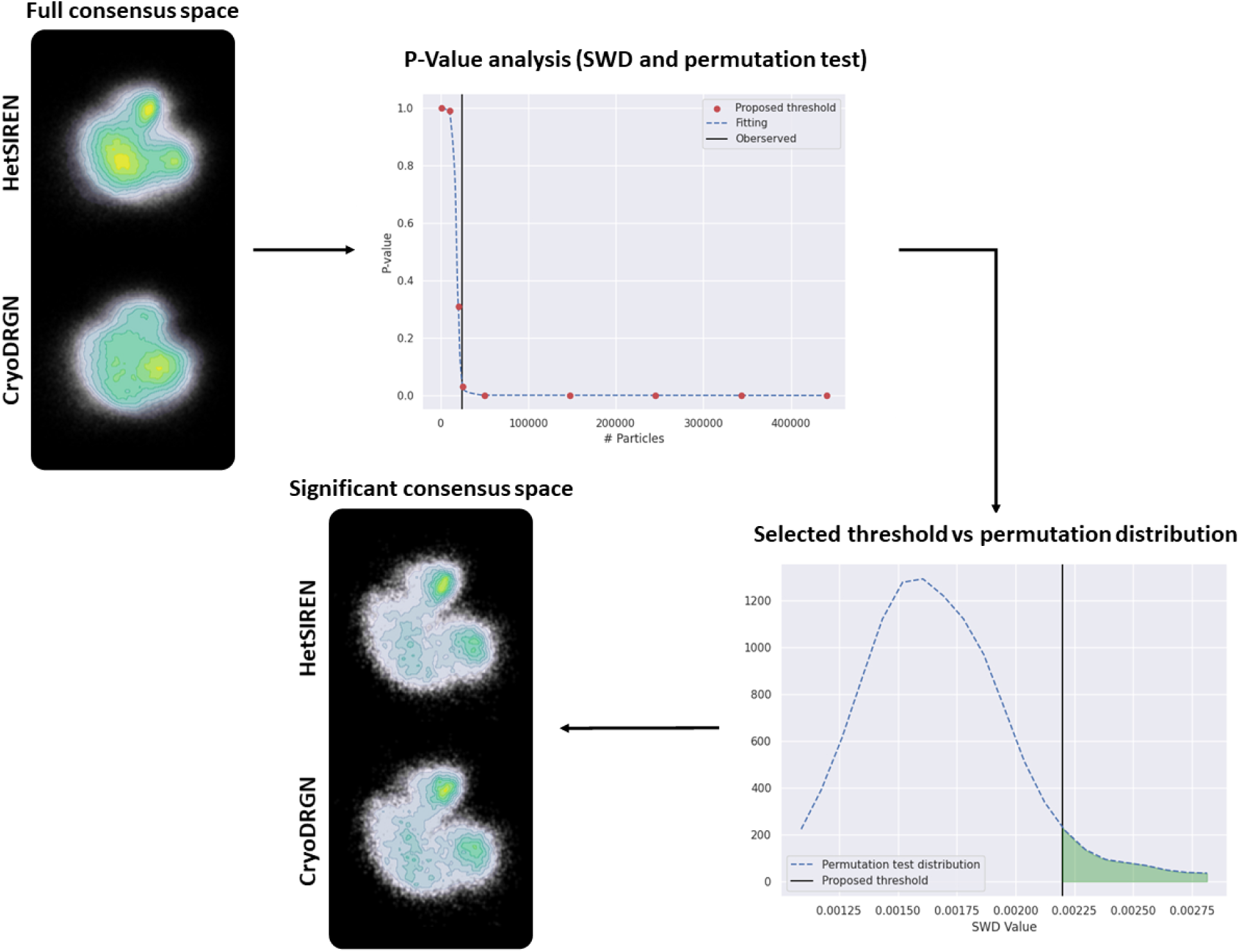
Scheme of the stabilization process to improve the state distribution reliability of the shared FlexConsensus process relies on the hypothesis that hetSIREN and CryoDRGN have found the same distribution al states. From this assumption, it is possible to determine the threshold veryfing the hypothesis, in a significant consensus space with a more accurate and reliable distribution of states.

## Discussion

The CryoEM community’s great interest in the emerging field of conformational variability analysis is reflected in the increasing number of advanced methods published in recent years. However, the interest in these tools raises a new challenge in comparing the estimations of different techniques to assess their stability, reliability, and accuracy. Moreover, the diverse nature of all these new methods makes comparing their results more complex, making it even harder to find robust approaches to accurately defining a valid consensus analysis.

To allow a better understanding of conformational landscapes, in this work, we discuss a new method to overcome the challenges arising from the comparison of different heterogeneity algorithms, leading to a whole range of possibilities to extract more reliable and accurate conformational states from independent conformational landscapes. Our new FlexConsensus approach relies on a multi-autoencoder architecture specifically designed to find the commonalities and differences of several conformational landscapes robustly, defining, in the last instance, a common consensus landscape with enhanced interpretability. From the consensus landscape, the network can automatically derive a consensus error metric for every particle in the conformational landscape, which can be used to assess the structural state’s reliability acroos heterogeneity methods explicitly determined for any given image in a dataset.

Thanks to FlexConsensus, filtering or cleaning a dataset based on its most stable conformational patterns can yield more robust subsets that can be confidently analyzed.

## Methods

This section describes the FlexConsensus multi-encoder architecture and the training strategy followed to allow the network to learn meaningful consensus spaces and consensus metrics from input conformational spaces.

### FlexConsensus multi-encoder architecture

FlexConsensus architecture consists of a multi-autoencoder architecture with a variable number of encoders and decoders determined by the total number of inputs. Each input corresponds to a conformational landscape that the consensus network is to consider, which may come from a different algorithm with its characteristics. A scheme of the architecture is provided in Figure 8.

**Figure 8:**
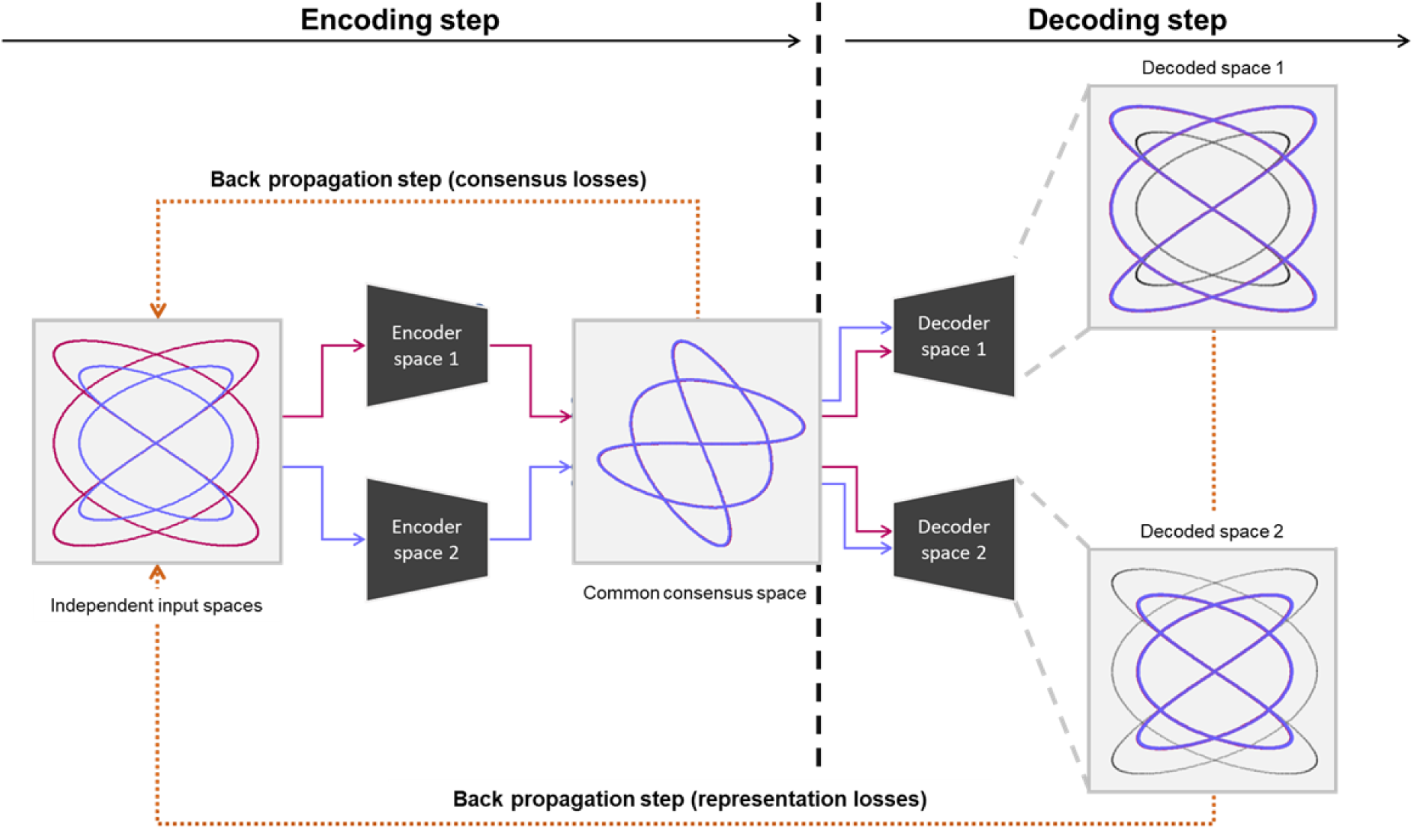
Scheme of FlexConsensus’ training workflow. A set of independent spaces is fed to different encoders, dependent representation of the initial spaces on a common consensus space. All encoded consensus forwarded to the first decoder, which is responsible for transforming the consensus spaces into a as possible to the first input space. A representation loss is evaluated and backpropagated through this step. The previous decoded step is repeated sequentially of all the decoders, allowing the he weights of the multi-encoder network.

Each input conformational landscape is forwarded through its encoder, which is composed of 3 fully connected layers with 1,024 neurons each and ReLU activation. The output layer of every encoder is then fed to a standard linearly activated latent space layer of a variable number of neurons that the user can choose. This bottleneck will become the common conformational space after the network is trained.

Lastly, each encoded data is forwarded through the corresponding decoder, trying to restore the original conformational space based on the information of the common latent space. Similarly to the encoders, each decoder comprises a set of 3 fully connected layers with 1,024 neurons each and ReLU activation, followed by a final layer with the same dimension as the corresponding initial conformational space and linear activation.

### Ensuring proper merging of conformational spaces

One of the main challenges in training a consensus network is correctly driving the network to merge spaces of varying characteristics in the same region of the latent space. Due to the properties of autoencoders, a simple mean-square error between the original and predicted spaces results in a latent space where different distributions are well separated. Although this will help decrease the network’s representation error, it is a highly undesirable effect, as it completely obscures the understanding of the intrinsic relations among the distributions of the input spaces.

A possible approach to solving the previous problem is enforcing the latent space distributions by ensuring their distances are as small as possible. This can be easily achieved by minimizing the mean-square error among all possible distribution pairs. In this way, the network is forced to learn a more meaningful representation compared to the original uncontiguous case.

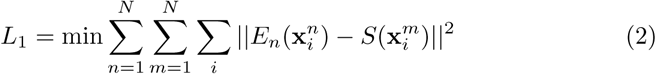

Being *N*, the total number of input spaces, and *E_n_*is the operator responsible for encoding a given space into the common consensus space.

However, the previous cost is not sufficient in our experiments, primarily due to the effect of the representation error of the decoder, which drove the result away from a fully merged consensus space. Therefore, an additional restriction is required to properly ensure that the local structure of the original conformational spaces is preserved as much as possible in the predicted consensus space. To that end, we proposed measuring the distances among the Shannon mappings of the original and predicted latent spaces. The Shannon mapping of every input and latent space batch is computed and posteriorly compared on a pairwise basis following a similar approach to the previous cost. The minimization of the all-to-all pairwise mean-square error among the Shannon mappings allows the network to learn to keep the original structure of the conformational spaces as consistent as possible. Therefore, the Shannon mapping cost function can be written as:

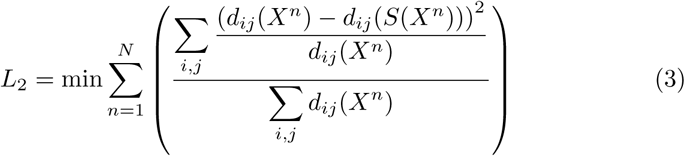

being *d*, the pairwise self-distance matrix computed from a set of coordinates, respectively, and *ɛ*, a small constant to prevent zero divisions. The self-distance matrix is computed as the all vs all distance among the coordinates in the set. In addition, a third cost is included in the total cost function to ensure that the distance distributions among the different consensus spaces are similar. To that end, we compute the self-pairwise distance matrix of the other encoded consensus spaces. The distance distribution is then calculated as the histogram of the self-pairwise distance matrices. Once the histograms are computed, they are compared pairwise by computing the Wasserstein distance between two different distributions, which is included as an extra regularization term in the cost function.

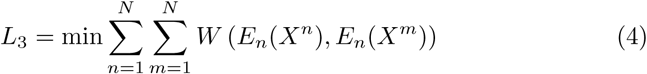

being *W* the function computing the Wasserstein distance between two sets of coordinates.

Combining the previous three costs at the latent space level in our experiments led to a good representation of a continuous consensus space. In addition to the last three cost functions, a standard representation error between the original and predicted conformational spaces based on the consensus space is also included. Since every conformational space might be affected by different estimation errors, only those regions consistently placed in the consensus space will have a low representation error. Therefore, the representation error provides a good measurement of the stability of the estimations, which can be used in conjunction with the consensus space to detect or filter out those regions in the conformational spaces that might have been estimated less reliably based on the agreement of all the spaces. The contribution of the previous regularizers in the overall cost function is summarized as follows:

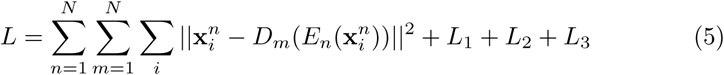

The first term represents the representation loss obtained by comparing the inputs against the decoded outputs, and *D_m_* is the corresponding decoder responsible for transforming a given consensus space into the input space *m*.

### FlexConsensus training strategy

The multi-encoder architecture proposed in FlexConensus introduces extra flexibility on the training step depending on the path to backpropagate the gradients computed during every forward pass through each autoencoder.

The workflow followed to train FlexConsensus is shown in Figure 8. In the simplified case proposed in the Figure, two independent input spaces are fed to their corresponding encoders, generating a different representation in a shared latent space. The first step includes encoding both input spaces, leading to two independent representations in the consensus space. Once the consensus spaces are generated, they are forwarded sequentially through the different decoders, triggering independent backpropagation steps for every decoder. We found the sequential training of the decoders to be more stable than training the whole network on a single backpropagation step, leading to a faster and more accurate convergence.

Once the network is trained, the corresponding encoder can recreate the complete consensus space and the location of a specific input in that space. In addition, the different encoder/decoder combinations can be used to convert to different input spaces and measure the consensus error estimated for a given input.

### Consensus set

Given the mapping of all points from the input spaces to a shared latent space, a natural question arises: which images indeed agree with one another? As illustrated in Fig. 1, we can calculate the distance between the projected points of Space 1 and their corresponding projections from Space 2 in the latent space. While this approach can be extended to more than two spaces, we simplify the explanation by focusing on two spaces. We then rank all points based on their average distance to points in the other space(s). Intuitively, the closest points represent agreement, while the most distant points indicate disagreement.

Using the top *N* particles from this list, we compare the distributions of their projections from Space 1 and Space 2. This comparison is based on the Wasserstein distance between the two sets of points. We approximate the Wasserstein distance in *k* dimensions to accelerate the calculations using the 1D Wasserstein distance of the points projected onto *κ* random unit vectors (13). We denote this observed distance as *d_obs_*. To assess its significance, we compare *d_obs_* to the distribution of distances obtained by randomizing the labels “Space 1” and “Space 2” (performing 100 randomizations). The proportion of randomized distances smaller than or equal to *d_obs_* gives the p-value, representing the probability of observing such a distance under random labeling. We expect that points with a small representation error in the latent space correspond to indistinguishable distributions of latent points from Space 1 and Space 2. However, as the number of particles *N* increases, a threshold is reached where the two distributions become clearly distinguishable, indicating that the two sets of points can no longer be considered equivalent. This threshold is defined as the point where the p-value falls below 0.05. The value of *N* at which this occurs establishes the set of consensus images.

## Data Availability

The experimental datasets analyzed in the manuscript are available in EMPIAR under the entries: 10028 [https://doi.org/10.6019/EMPIAR-10028]. The e SARS-CoV-2 D614 spike dataset will be published in a different work in the future.

## Code Availability

The FlexConsensus algorithm is available through Scipion 3.0 (10) under the plugins and scipion-em-flexutils. The protocol corresponding to the algorithm described in this manuscript is flexutils - train - FlexConsensus and flexutils - interactive consensus - FlexConsensus.

## Author Contributions

DH developed and tested the FlexConsensus method presented throughout the manuscript. CPM helped with the data preprocessing. COSS and JMC jointly supervised this work.

## Competing Interests

The authors declare that they have no competing interests.

## Acknowledgments

The authors acknowledge the financial support from the Ministry of Science, Innovation and Universities (BDNS n. 716450) to the Instruct Image Processing Center (I2PC) as part of the Spanish participation in Instruct-ERIC, the European Strategic Infrastructure Project (ESFRI) in the area of Structural Biology, Grant [PID2022-136594NB-I00] funded by MICIU /AEI/10.13039/501100011033/ and “ERDF A way of making Europe”, by the “European Union” Spanish State Research Agency, AEI/10.13039/501100011033, through the “Severo Ochoa” Programme for Centres of Excellence in R&D [CEX2023-001386-S] Comunidad Autónoma de Madrid” through Grant: S2022/BMD-7232, European Union (EU) and Horizon 2020 through grant: HighResCells (ERC - 2018 - SyG, Proposal: 810057) European Union (EU), and Horizon Europe through grant: Fragment Screen Proposal: 101094131. We thank Professor S. Subramaniam for providing us with the data processed and presented in this work, as well as for his support and feedback.

---

**Supplementary Figure 1:**
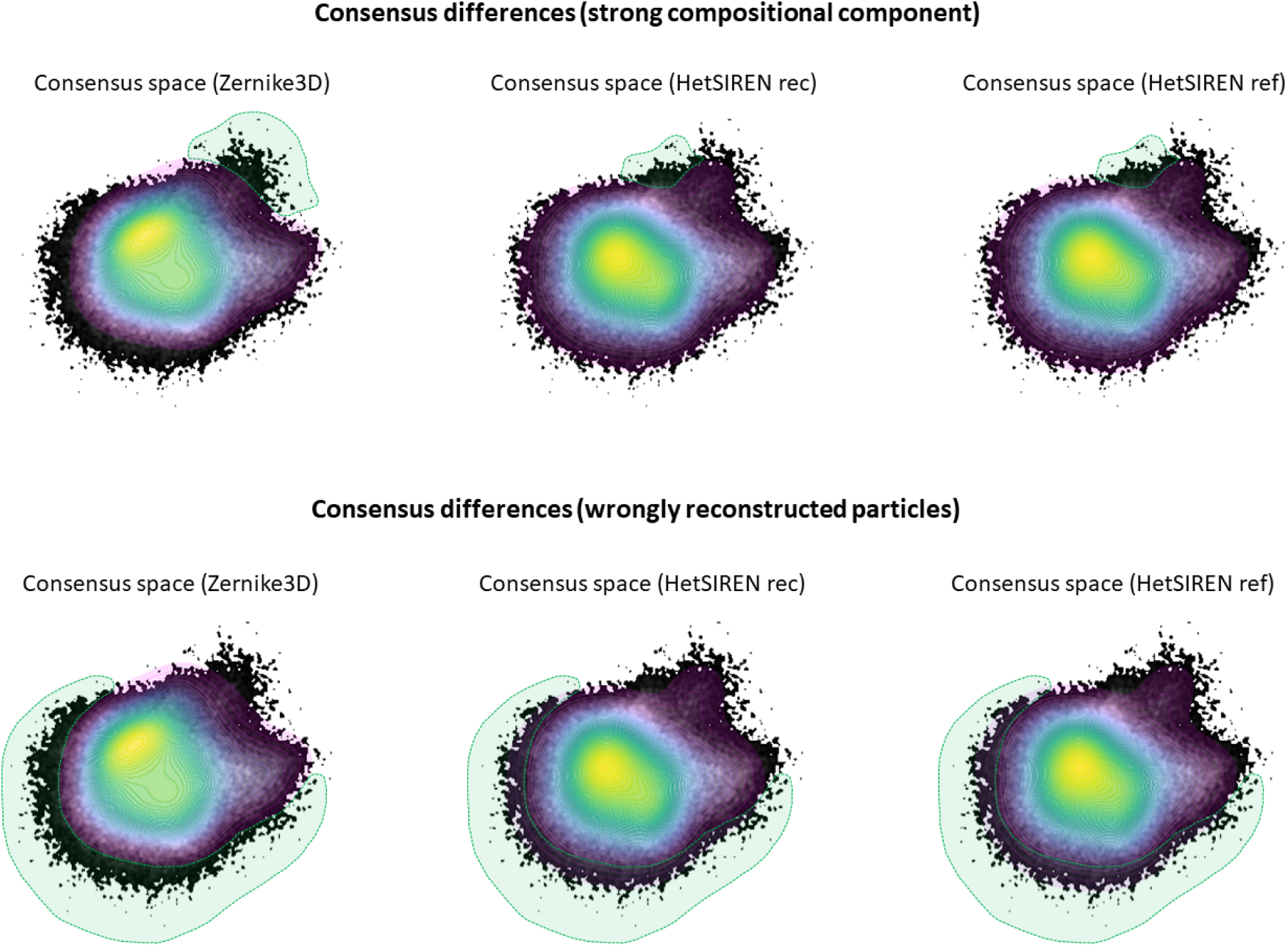
Highlight of the main differences found in the consensus landscape estimated for the dataset. The top panel shows the regions where Zernike3D and HetSIREN disagree due to the trong compositional change that Zernike3D cannot capture. The bottom panel shows a strong e periphery of the landscape, mainly associated with wrongly estimated conformations with high e of HetSIREN. Since Zernike3D does not have the capacity to add noise to the conformation it riphery is closer to the central region of the consensus space, as shown in the figure.

## Notes

### Competing Interest Statement

The authors have declared no competing interest.

